# vcferr: Development, Validation, and Application of a SNP Genotyping Error Simulation Framework

**DOI:** 10.1101/2022.03.28.485853

**Authors:** V. P. Nagraj, Matthew Scholz, Shakeel Jessa, Jianye Ge, August E. Woerner, Bruce Budowle, Meng Huang, Stephen D. Turner

**Affiliations:** Signature Science, LLC., Austin, TX 78759, USA; Center for Human Identification, Department of Microbiology, Immunology, and Genetics, University of North Texas Health Science Center, Fort Worth, TX, 76107, USA

**Keywords:** bioinformatics, kinship, python, genealogy, gwas, simulation, benchmarking

## Abstract

**Motivation:** Genotyping error can impact downstream SNP-based analyses. Simulating various modes and and levels of error can help investigators better understand potential biases caused by miscalled genotypes.

**Results:** We have developed and validated `vcferr`, a tool to probabilistically simulate genotyping error and missigness in VCF files. We demonstrate how `vcferr` could be used to address a research question by introducing varying levels of error of different type into a sample in a simulated pedigree, and assessed how kinship analysis degrades as a function of kind and type of error.

**Software Availability:** `vcferr` is available for installation via PyPi (https://pypi.org/project/vcferr/) or conda (https://anaconda.org/bioconda/vcferr). The software is released under the MIT license with source code available on GitHub (https://github.com/signaturescience/vcferr).

## Introduction

Single nucleotide polymorphisms (SNPs) are inherited single base-pair substitutions in genomic DNA that are easily characterized by microarray, targeted, or whole-genome sequencing, and have a wide range of uses in personalized medicine, disease association studies, population genetics, genealogy, and forensics [1, 2]. Genotyping error occurs at known rates on genomic sequencing and array platforms [3]. However, novel applications of sequencing or genotyping such as those encountered in forensics [4] or ancient DNA [5], or novel downstream data analysis techniques such as imputation on low-pass whole-genome sequencing [6, 7, 8] can result in genotyping errors that differ from well-established rates and types of errors that occur in typical genotyping and high-coverage sequencing in high-quality samples. Different applications or data analysis procedures can result in errors that can manifest in a variety of ways, for example, with differing rates of drop in or drop out of heterozygous calls.

There are several existing tools for simulating sequencing or genotype array data. Wgsim is a tool for simulating sequencing reads from a reference genome, allowing for a uniform user-defined substitution sequencing error [9]. The ART sequencing read simulator allows for technology-specific read error models and base quality value profiles derived empirically from large sequencing datasets [10]. Downstream of raw sequencing read error simulation, other tools provide methods for simulating genotyping data (as if variant calling had already been performed, or genotyping microarrays were being used to generate SNP genotypes). The pedsim software allows for simulating large, arbitrarily complex pedigrees with genotypes drawn from founders in a VCF [11], allowing for realistic familial data to be generated by employing sex-specific genetic maps and realistic crossover interference models [12]. Ped-sim allows for specifying the rate of genotyping error and the rate of an opposite homozygous error conditional on a genotyping error at a marker, but does not allow for arbitrary specification of substitution rates in biallelic SNPs (such as different rates for heterozygous to homozygous reference drop-out versus homozygous reference to heterozygous drop-in), and it applies these universal error rates to all samples in a multi-sample VCF.

While there are tools available to simulate errors in *sequencing reads*, and other tools such as ped-sim which allow for broad specification of global error rate in *genotype calls*, to our knowledge there is no existing tool that allows for individual specification of the type and degree of genotyping error an arbitrary sample in a VCF (e.g., different rates for heterozygous to homozygous reference versus homozygous to heterozygous). The ability to precisely simulate the *degree* and *kind* of genotyping error (based on error probabilities discovered from prior upstream data analysis) is necessary to facilitate testing and evaluation of downstream applications which may be differentially sensitive to different types of genotyping errors. Here we describe our efforts to develop, validate, and demonstrate an application and usage of `vcferr`, a user-friendly python tool for probabilistically simulating arbitrarily specified genotyping error and missing data into VCF files. Source code, PyPI/bioconda package, and code to reproduce the results presented here are given in the Software Availability section below.

## Methods

### Implementation

#### The vcferr python package

`vcferr` is written in Python and distributed to run directly from a command line interface. The tool is built using `pysam` [13], which internally manages reading and writing VCF file contents. When run on an input VCF, `vcferr` will load all variant records for a given sample. The tool checks each genotype, and randomly draws from a list of possible genotypes (heterozygous, homozygous for the alternate allele, homozygous for the reference allele, missing) with each element weighted by error rates specified by the user. The processing runs iteratively for every site in the input VCF, with the output streamed or optionally written to a new output VCF file. The following error and missingness models are available:

- `rarr` = Heterozygous drop out to homozygous reference: `0/1` or `1/0` to `0/0`
- `aara` = Homozygous alternate to heterozygous: `1/1` to `0/1`
- `rrra` = Homozygous reference to heterozygous drop in: `0/0` to `0/1`
- `raaa` = Heterozygous to homozygous alternate: `0/1` or `1/0` to `1/1`
- `aarr` = Homozygous alternate to homozygous reference: `1/1` to `0/0`
- `rraa` = Homozygous reference to homozygous alternate: `0/0` to `1/1`
- `ramm` = Heterozygous to missing: `0/1` or `1/0` to ./.
- `rrmm` = Homozygous reference to missing: `0/0` to ./.
- `aamm` = Homozygous alternate to missing: `1/1` to ./.

We validated `vcferr` by reviewing concordance metrics between VCF files before and after error simulation. To perform this validation we used data from the 1000 Genomes Project [14] and the companion error assessment tool described below. First, we retrieved all genotype calls for chromosome 22 from the 1000 Genomes FTP site. Next, we simulated a set of error rates for one sample. We observed that the proportion of mismatches reported by our `bcftools`-based error assessment procedure aligned as expected to each of the specified error probabilities. We also confirmed that the simulation of error in one sample did not trigger any mismatches for other samples in the VCF. The chromosome 22 VCF includes 1,103,547 positions and 2504 samples, and genotype error simulation took ∼5 minutes on a single CPU. The code used to fully reproduce this validation analysis is described in the Software Availability section below.

### Companion evaluation tool: nrc

As part of this analysis we developed a companion tool (called `nrc`) to measure all types of error types that can be observed with biallelic SNPs between a reference and a query sample. This tool also computes the nonreference concordance (NRC), which is defined as:

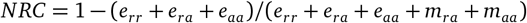

Where *e*_*rr*_, *e*_*ra*_, and *e*_*aa*_, are defined as the the counts of mismatches for the homozygous reference, heterozygous, and homozygous ALT genotypes, and *m*_*ra*_, and *m*_*aa*_ are the counts of the matches at the heterozygous and homozygous ALT genotypes (the number of homozygous reference matches, *m*_*rr*_, is omitted from this analysis). We implemented this procedure as a Docker container, which first runs `bcftools stats` [13] followed by post-processing results using R. The Dockerfile needed to build the `nrc` image is available under the MIT license at https://github.com/signaturescience/nrc, and the Docker image can be pulled directly from the GitHub container registry with docker pull ghcr.io/signaturescience/nrc.

### Operation

`vcferr` is packaged in Python and available to be installed from PyPi or Bioconda. Once installed, the tool can be used through the command line without calling the Python interpreter. `vcferr` requires a path to an input VCF from which error will be simulated as the first positional argument. The tool also requires a specification for `--sample` for the ID of the sample to be simulated as it appears in the VCF header. All other error flags are optional, as is `--output-vcf` which if used will define the path to which the VCF should be written. If no value is passed to a given error or missingness option, then the probability will default to 0. The following is a minimal example of usage, which will result in 5% of the heterozygous genotypes in “sampleX” to be erroneously called as homozygous reference, and 1% of that same individual’s heterozygous genotypes to be erroneously called as homozygous alternate, with output being written to a new bgzipped VCF file:

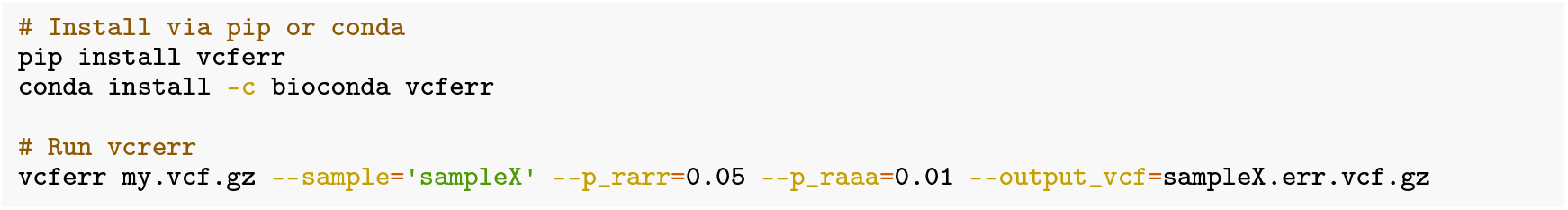

The companion tool `nrc` for assessing individual error rate probabilities and overall non-reference concordance is available as a Docker image from the GitHub container registry. Once installed, the container can be run through the command line with no dependencies other than Docker. The container expects two VCF files and a sample ID which exists in both VCFs to compare. Optionally, a set of sites at which to compare can be specified (default is to compare all sites). This is useful to restrict analyses to a subset of sites (e.g., those that have at least some minimum minor allele frequency from GnomAD, given that filtered site VCF). Example usage:

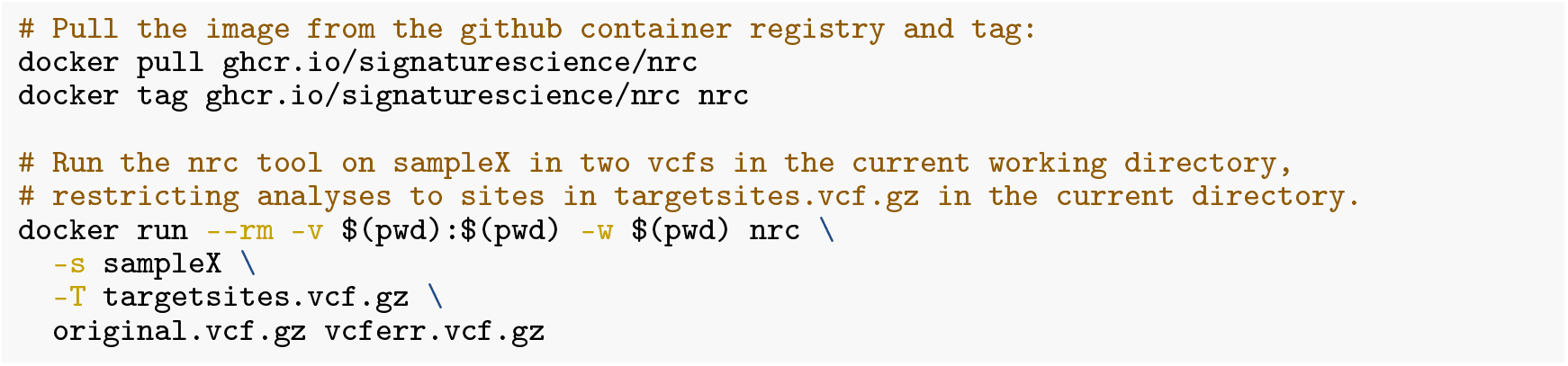

## Use Cases

### Benchmarking SNP-based kinship analysis

To demonstrate how `vcferr` could be used to address a research question, we conducted an analysis of the impact of genotyping error on pairwise kinship estimation. For instance, a heterozygous drop-out to homozygous reference may be much better tolerated than a switch from homozygous reference to homozygous alternate. We first generated simulated pedigree data using `ped-sim` [12] with founders drawn from the 1000 Genomes GBR population. The simulated data was masked to a subset of 590,241 sites, including content from the Illumina Global Screening Array. With all data originally simulated to include no error, we next identified an individual for which we would inject genotyping error. We used `vcferr` to iterate through each error mode from 0 to 20% error at 1% steps, holding all other error modes constant at 0.

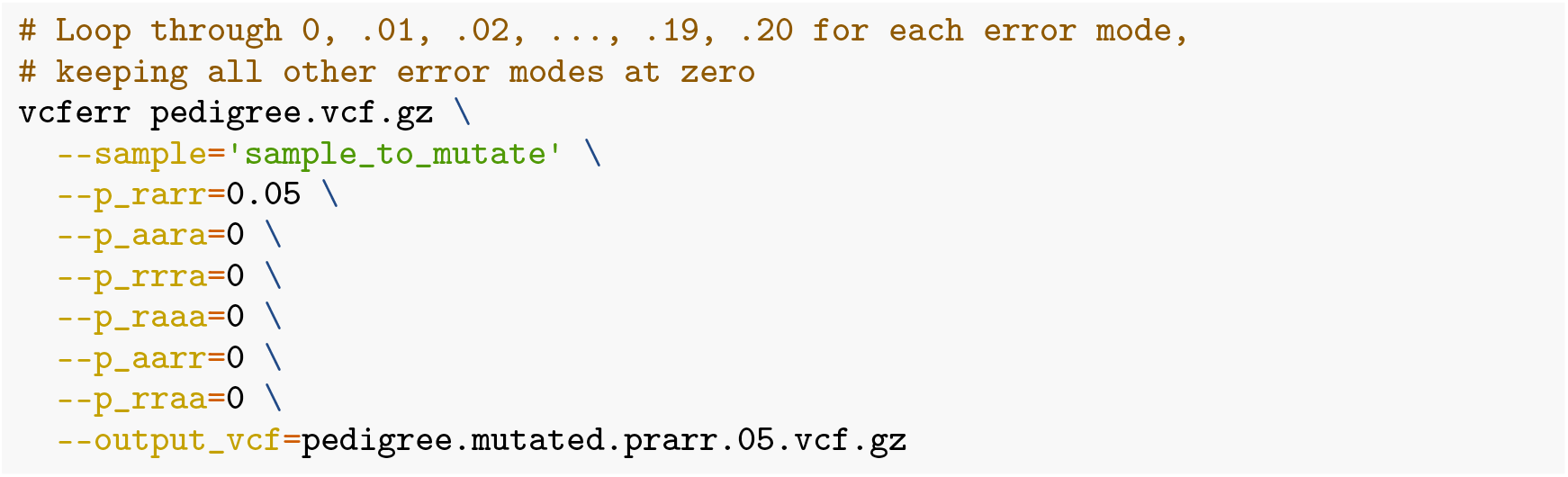

Next, we obtained KING robust [15] kinship coefficients from `akt` [16] for all pairwise comparisons that included the error-simulated individual in the resulting simulation VCFs. We then used the kinship coefficients to infer relatedness out to the third-degree and compared accuracy of degree inference to truth from the original simulated pedigree. The results of this analysis are summarized in Figure 1. Code needed to reproduce this entire analysis, implemented as a Snakemake workflow [17], is available in the Zenodo archive described in the Software Availability section below.

**Figure 1.**
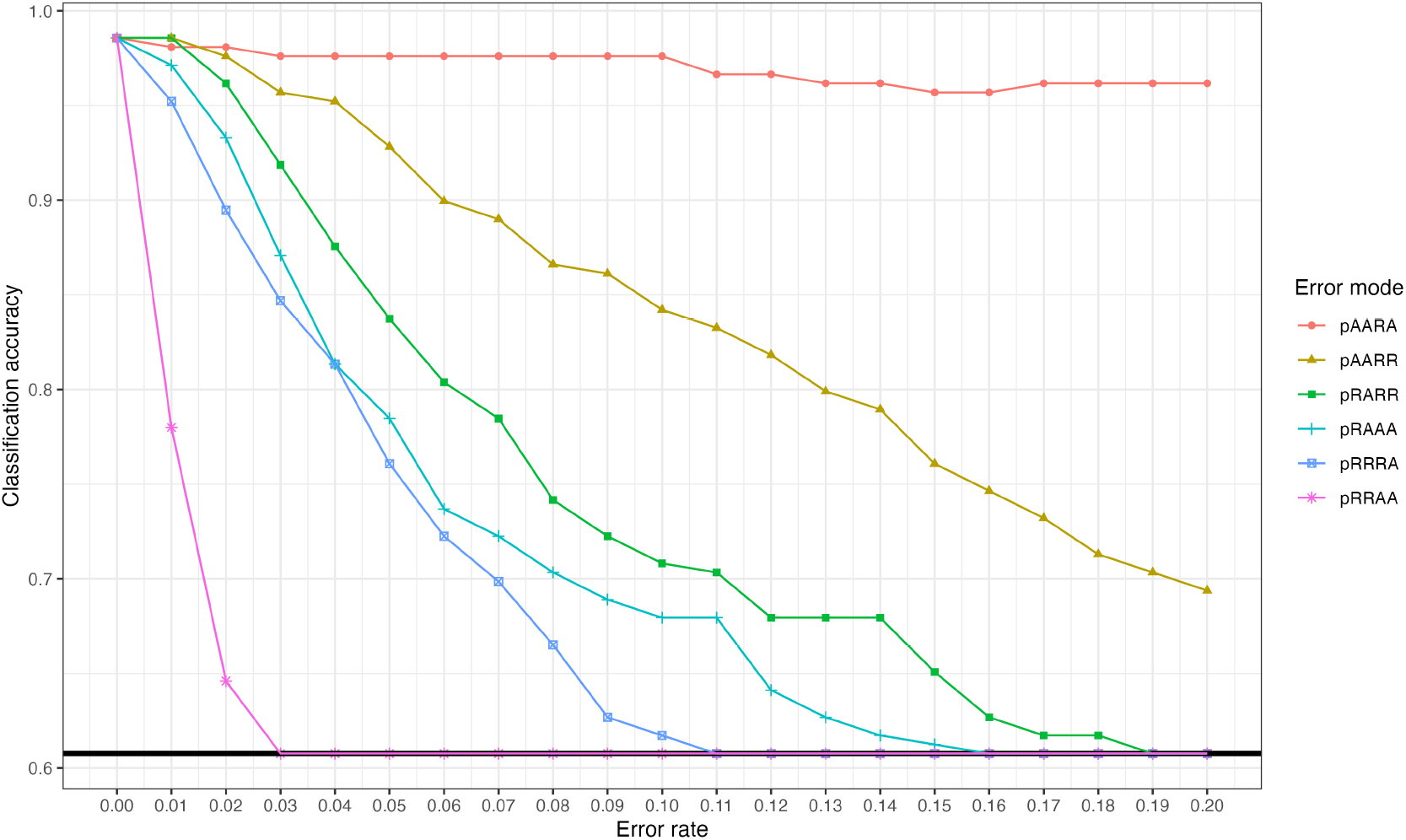
Impact of genotyping error on inferred degree classification accuracy. Each line corresponds to a type of error, with error rates from 0 to 0.2 stepped at 0.01 increments while holding all other types of error constant at 0. The solid black line at bottom represents the accuracy at guessing, which is the floor for relationship degree classification.

In short, certain kinds of error dramatically decrease accuracy of relatedness inference with the KING robust estimator. Note that the sample for which errors were simulated included 430,056 homozygous reference sites, 105,413 heterozygous sites, and 54,772 homozygous alternate site, resulting in varying absolute counts of errors being simulated at each 0 to 20% step (as expected in any sample with genotypes from microarray or sequencing). By leveraging `vcferr` to inject a range of genotyping error scenarios, we can identify the scenarios that are most likely to impact kinship analysis.

### Genome-wide association studies and polygenic risk scores

Genome-wide association studies (GWAS) are a widely used approach for genetic epidemiology studies attempting to understand genetic underpinnings to complex human disease [18]. Formalin-fixed, paraffin-embedded (FFPE) tissue fixation is a standard means to preserve tissues from clinical samples, and can result in genotyping error [19] that can vary by type, with heterozygous errors being more commonly discordant [20]. Power and sample size requirements for GWAS are impacted by genotyping error [21]. If error probabilities for different sample types such as FFPE tissue samples are known, the impact of different types of genotyping error on GWAS power or sample size requirements can be estimated in simulation. One of many available tools can be used to simulate GWAS data with known sample size, effect size, SNP density, etc. [22, 23]. Following data simulation, vcferr can be used to inject error in samples of interest, precisely controlling the degree and kind of error in each sample.

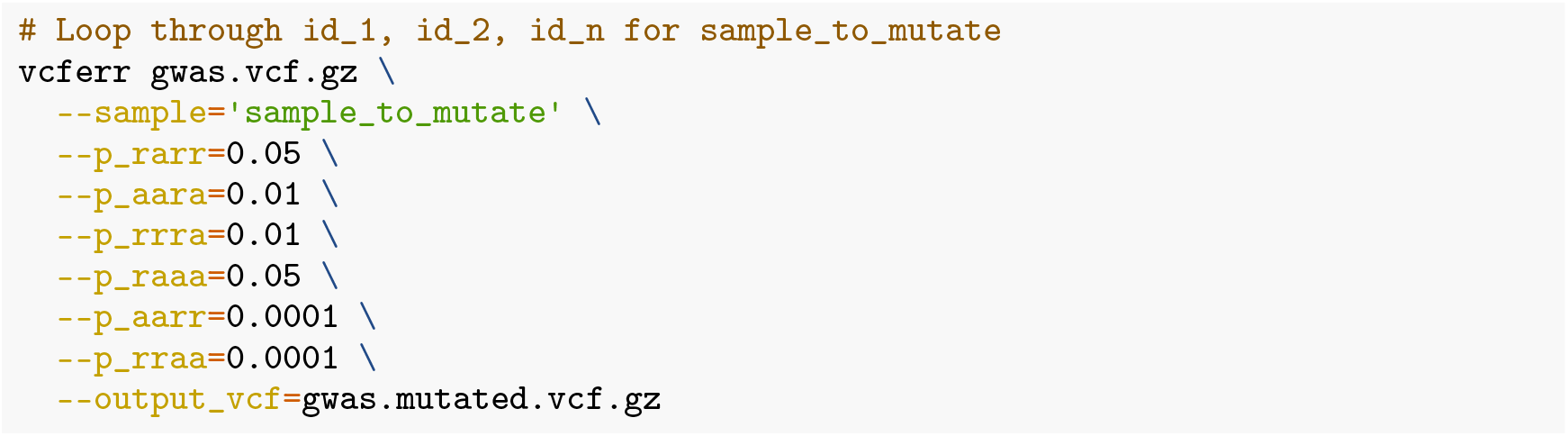

Similarly, a polygenic risk score (PRS) captures the additive effect of hundreds, or perhaps thousands of SNPs, and are computed on genome-wide genotyping data using a prediction model based on summary statistics from external GWAS studies [24]. PRS are first calculated by summing the risk alleles an individual has where each risk allele is weighted by the risk allele effect size as estimated by a large GWAS on that phenotype. The PRS is then applied to target genotype data to assess overall genetic liability to developing the trait or disease. If the target genotypes result from a process that introduces error at different levels for different zygosity (e.g., low-pass whole genome sequencing), the performance of a PRS will be impacted differently depending on the type and kind of error introduced. If this error profile is known by characterizing against high-quality samples, those error profiles can be used to inject error at those specified rates into samples simulated for assessing the predictive ability of a PRS in a new target genotype collection using the same approach above for GWAS.

## Summary

`vcferr` provides a flexible, user-friendly interface to simulate genotyping error in VCF files. While tools exist to simulate entire VCFs with error (e.g. `ped-sim` [12]), to our knowledge at the time of writing there are currently no other tools that can simulate arbitrary individuals in the VCF at varying types and levels of error while leaving all other samples as-is. We anticipate that `vcferr` will be useful for researchers in multiple areas, including forensic genomics and medical genetics.

## Software availability

### vcferr

The `vcferr` python tool is the primary software tool described and demonstrated in this paper.

1. Software available from: PyPI (https://pypi.org/project/vcferr/) and Bioconda (https://anaconda.org/bioconda/vcferr).
2. Source code available from: https://github.com/signaturescience/vcferr.
3. Archived source code at time of publication: https://doi.org/10.5281/zenodo.6611796.
4. Software license: MIT License.

### nrc

The `nrc` companion tool was created for assessing error produced by `vcferr`.

1. Software (Docker image) available from: https://ghcr.io/signaturescience/nrc.
2. Source code available from: https://github.com/signaturescience/nrc.
3. Archived source code at time of publication: https://doi.org/10.5281/zenodo.6611782.
4. Software license: MIT License.

### Code for reproducing the analyses in this paper

Code for reproducing the analyses presented in this paper is available and documented on Zenodo at https://doi.org/10.5281/zenodo.6599133.

## Author information

VPN, SDT, and MBS developed the software.

All authors contributed to method development. VPN wrote the first draft of the manuscript.

All authors assisted with manuscript revision.

All authors read and approved the final manuscript.

## Competing interests

No competing interests were disclosed.

## Grant information

This work was supported in part by award 2019-DU-BX-0046 (Dense DNA Data for Enhanced Missing Persons Identification) to B.B., awarded by the National Institute of Justice, Office of Justice Programs, U.S. Department of Justice and by internal funds from the Center for Human Identification. The opinions, findings, and conclusions or recommendations expressed are those of the authors and do not necessarily reflect those of the U.S. Department of Justice.

## Acknowledgments

The authors acknowledge Dr. Mike Coble for helpful discussions about the software and its use cases.

## Notes

https://github.com/signaturescience/vcferr/

https://pypi.org/project/vcferr/

https://anaconda.org/bioconda/vcferr

https://github.com/signaturescience/nrc

### Competing Interest Statement

The authors have declared no competing interest.

### Summary of Updates

Bring information from the supplement into the main text; expand use cases; format for F1000Research submission.

