## Supplementary Information for "vcferr: Development, Validation, and Application of a SNP Genotyping Error Simulation Framework"

### Documentation for reproducing analyses presented in: `vcferr`: Development, Validation, and Application of a SNP Genotyping Error Simulation Framework

### 1 Validation analyis

#### 1.1 Summary

The code below provides steps necessary to conduct the validation analysis. Besides `vcferr` the original analysis used the following software (with versions):

- `bcftools` 1.9
- `tabix` 1.9

Both can be installed with `conda create  -c bioconda -n bcftools bcftools=1.9 tabix=1.9`.

Note that comparison procedure uses `nrc`, which is distributed as a Docker image and can be built using code and instructions at https://github.com/signaturescience/nrc.

#### 1.2 Prep and analysis pipeline

```
WORKDIR=/path/to/working/directory

wget -P $WORKDIR \
    http://ftp.1000genomes.ebi.ac.uk/vol1/ftp/release/20130502/ALL.chr22.phase3_shapeit2_mvncall_integrated_v5b.20130502.genotypes.vcf.gz
wget -P $WORKDIR \
    http://ftp.1000genomes.ebi.ac.uk/vol1/ftp/release/20130502/ALL.chr22.phase3_shapeit2_mvncall_integrated_v5b.20130502.genotypes.vcf.gz.tbi

vcferr $WORKDIR/ALL.chr22.phase3_shapeit2_mvncall_integrated_v5b.20130502.genotypes.vcf.gz \
    --sample 'NA20808' \
    --p_rarr=0.02 \
    --p_aara=0.09 \
    --p_rrra=0.08 \
    --p_aarr=0.05 \
    --p_rraa=0.07 \
    --p_raaa=0.1 \
    --output_vcf=$WORKDIR/1kg.NA20808.chr22.all.vcf.gz

tabix $WORKDIR/1kg.NA20808.chr22.all.vcf.gz

bcftools view -s NA20808 $WORKDIR/ALL.chr22.phase3_shapeit2_mvncall_integrated_v5b.20130502.genotypes.vcf.gz -Oz -o $WORKDIR/NA20808.chr22.vcf.gz
tabix $WORKDIR/NA20808.chr22.vcf.gz

docker run --rm -v $WORKDIR:/data nrc:latest /data/NA20808.chr22.vcf.gz /data/1kg.NA20808.chr22.all.vcf.gz > 1kg.NA20808.chr22.all.nrc

bcftools view -s NA21144 $WORKDIR/ALL.chr22.phase3_shapeit2_mvncall_integrated_v5b.20130502.genotypes.vcf.gz -Oz -o $WORKDIR/NA21144.chr22.vcf.gz
tabix $WORKDIR/NA21144.chr22.vcf.gz

docker run --rm -v $WORKDIR:/data nrc:latest /data/NA21144.chr22.vcf.gz /data/1kg.NA20808.chr22.all.vcf.gz > 1kg.NA21144.chr22.all.nrc
```

### 2 Demonstration analysis

#### 2.1 Summary

The code below provides steps necessary to conduct the demonstration analysis. Besides `vcferr` the original analysis used the following software (with versions):

- `bcftools` 1.9
- `tabix` 1.9
- `akt` 0.3.3
- `ped-sim` 1.1.17
- `snakemake` 3.13.3

To run the analysis, the directory should be organized as follows:

```
├── figure1.R
├── Snakefile
└── data
    ├── input
    │   ├── gen.sh
    │   ├── GBR.txt
    │   └── 5GHS.def
    ├── akt
    └── vcferr
```

Begin by running `gen.sh` to retrieve and prepare 1000 Genomes data and simulate a pedigree with founders from GBR population using `ped-sim`. The data generation step uses the `5GHS.def` file and `GBR.txt` for pedigree definition and 1000 Genomes samples. Note that the retrieval and concatenation of autosomal data from 1000 Genomes is the longest step in processing, and may take hours to complete depending on computational resources available.

Once the `ped-sim` data generation step is complete, run the `snakemake` pipeline defined in `Snakefile` to inject genotyping error at various rates with `vcferr` and estimate pairwise kinship coefficients with `akt`. Tool-specific result files will be written to the respective directories in the `data/` directory.

To recreate the figure run `figure1.R`, which includes code to read in the `akt` results, resolve inferred degrees from kinship coefficients, and compare accuracy of degree inference to true relatedness per the original simulated pedigree.

Annotated contents of the files mentioned above are provided below.

#### 2.2 Data generation pipeline (gen.sh)

```
# First ...
# 1. Download all 1000G from ftp://ftp.1000genomes.ebi.ac.uk/vol1/ftp/release/20130502/
# 2. Concatenate all autosomal biallelic SNPs into a single vcf.gz
# 3. Subset concatenated chromosomes to only sites that are on a microarray platform

# wget the entire 20130502 release variant calls and all associated data
wget -r --level=1 ftp://ftp.1000genomes.ebi.ac.uk/vol1/ftp/release/20130502/
# Move it all into a folder called 'ftp'
mkdir -p ftp
mv ftp.1000genomes.ebi.ac.uk/vol1/ftp/release/20130502/* ftp/
rm -rf ftp.1000genomes.ebi.ac.uk/

# CONCATENATE ALL AUTOSOMES
# /tmp is too small to perform the sort. Make a tmp dir here.
mkdir -p tmp
# concat all autosomes, get only biallelic snps, sort, write out to file, index (takes most of a day)
bcftools concat -a ftp/ALL.chr[0-9]*.vcf.gz | bcftools view -v snps -m2 -M2 | bcftools sort --temp-dir ./tmp/ -Oz -o 1000g.concat.autosomes.vcf.gz
tabix 1000g.concat.autosomes.vcf.gz

# Next ...
# 1. Download, unzip, and dos2unix format the Illumina GSA manifest
# 2. Create a tab-delimited file ready for bcftools --regions-file

# Get the manifest directly from Illumina, unzip, remove incompatible line endings, clean up
wget https://support.illumina.com/content/dam/illumina-support/documents/downloads/productfiles/global-screening-array-24/v3-0/GSA-24v3-0-A1-manifest-file-csv.zip
unzip GSA-24v3-0-A1-manifest-file-csv.zip
dos2unix GSA-24v3-0_A1.csv
rm GSA-24v3-0-A1-manifest-file-csv.zip

# Parse that manifest
# The last 20 or so lines are unformatted "control" sequences. The sed line gets rid of everything including and after a line that contains "Controls"
# awk: F, indicates fields separated by commas; OFS="\t" indicates output should be tab delimited. NR>8 skips the first 8 line header, columns 10 and 11 contain chr/position.
# grep out autosomes, sort numerically and get unique rows
cat GSA-24v3-0_A1.csv | sed '/Controls/Q' | awk -F, -v OFS="\t" 'NR>8 {print $10,$11}' | grep "^[1-9]" | sort -nk1,2 | uniq > GSA-24v3-0_A1.csv.autosomes.chr-pos.tsv

# Get the "reliable" sites from the AKT pruned sites from https://github.com/Illumina/akt/tree/master/data
wget https://raw.githubusercontent.com/Illumina/akt/master/data/wgs.grch37.vcf.gz -O reliable.grch37.vcf.gz
wget https://raw.githubusercontent.com/Illumina/akt/master/data/wgs.grch37.vcf.gz.tbi -O reliable.grch37.vcf.gz.tbi
bcftools query -f '%CHROM\t%POS\n' reliable.grch37.vcf.gz > reliable.grch37.chr-pos.tsv

# Concatenate GSA autosomes and reliable sites, get autosomes, sort, write to tsv for bcftools view -R
cat GSA-24v3-0_A1.csv.autosomes.chr-pos.tsv reliable.grch37.chr-pos.tsv | grep "^[1-9]" | sort -nk1,2 | uniq  > GSA-reliable.chr-pos.tsv

# Next ...
# 1. Download the sex-specific recombination
# 2. Assemble map file for ped-sim

# per ped-sim documentation https://github.com/williamslab/ped-sim#map-file
wget https://github.com/cbherer/Bherer_etal_SexualDimorphismRecombination/raw/master/Refined_genetic_map_b37.tar.gz
tar xvzf Refined_genetic_map_b37.tar.gz
printf "#chr\tpos\tmale_cM\tfemale_cM\n" > refined_mf.simmap
for chr in {1..22}; do
  paste Refined_genetic_map_b37/male_chr$chr.txt Refined_genetic_map_b37/female_chr$chr.txt \
    | awk -v OFS="\t" 'NR > 1 && $2 == $6 {print $1,$2,$4,$8}' \
    | sed 's/^chr//' >> refined_mf.simmap;
```

```
done

# clean up
rm -rf Refined_genetic_map_b37*

# Next ...
# 1. Get ped-sim interference file
wget https://raw.githubusercontent.com/williamslab/ped-sim/master/interfere/nu_p_campbell.tsv

# Next ...
# 1. Subset the autosomal 1000G data to GSA+reliable sites for unreleated GBR samples
# 2. Index the output file
# NOTE: the contents of GBR.txt includes sample IDs for all unrelated GBR samples
bcftools view -R GSA-reliable.chr-pos.tsv 1000g.concat.autosomes.vcf.gz -S GBR.txt -v snps -Oz | bcftools sort -Oz -o 1000g.concat.autosomes.GBR.vcf.gz
tabix 1000g.concat.autosomes.GBR.vcf.gz

# Next ...
# 1. Run ped-sim
# 2. Zip and index result file
# NOTE: the contents of the pedigree definition includes 5 generations with half-siblings

ped-sim \
    --seed 090322 \
    --nogz \
    --err_rate 0 \
    --miss_rate 0 \
    --err_hom_rate 0 \
    -d 5GHS.def \
    -m refined_mf.simmap \
    --intf nu_p_campbell.tsv \
    --fam \
    -i 1000g.concat.autosomes.GBR.vcf.gz \
    -o simulation

bgzip simulation.vcf
tabix simulation.vcf.gz
```

#### 2.3 Contents of pedigree definition file (**5GHS.def**)

```
# Make lots of cousins.
# some branches will have half-siblings
# 1 copy, 5 generations
######
# half_siblings requires 18 founders/copy
####
def halfSiblings 1 5
1 1
# 2nd generation has +1 offspring that does not form branch (+2 form new branches)
2 2 8
# 3rd generation has +1 offspring that does not branch, 2 additional branches formed, for a total of 4
# branch 1 and branch 2 come from 1 same parent, different spouse
3 4 8 1:1 2:1 3-4:2
#same, doubling
# only making 1/2 siblings for branches 1 and 2
4 4 16 1:1 2:1 3-4:2 5-6:3 7-8:4
#each of 8 branches has 2 offspring.
5 4
```

#### 2.4 Contents of sample list file (GBR.txt)

```
HG00096
HG00097
HG00099
HG00100
HG00101
HG00102
HG00103
HG00105
HG00106
HG00107
HG00108
HG00109
HG00110
HG00111
HG00112
HG00113
HG00114
HG00115
HG00116
HG00117
HG00118
HG00119
HG00120
HG00121
HG00122
HG00123
HG00125
HG00126
HG00127
HG00128
HG00129
HG00130
HG00131
HG00132
HG00133
HG00136
HG00137
HG00138
HG00139
HG00140
HG00141
HG00142
HG00143
HG00145
HG00146
HG00148
HG00149
HG00150
HG00151
HG00154
HG00155
HG00157
HG00158
HG00159
HG00160
HG00231
HG00232
HG00233
HG00234
HG00235
HG00236
HG00237
HG00238
HG00239
HG00240
HG00242
HG00243
HG00244
HG00245
HG00246
HG00250
HG00251
HG00252
HG00253
HG00254
HG00255
HG00256
HG00257
HG00258
HG00259
HG00260
HG00261
HG00262
HG00263
HG00264
HG00265
HG01334
HG01789
HG01790
HG01791
HG02215
```

#### 2.5 Snakemake instructions to run `vcferr` and `akt` (`Snakefile`)

```
###############

import glob as glob
import os
import re
import numpy as np

PRARR=[0,0.01,0.02,0.03,0.04,0.05,0.06,0.07,0.08,0.09,0.1,0.11,0.12,0.13,0.14,0.15,0.16,0.17,0.18,0.19,0.2]
PAARA=[0,0.01,0.02,0.03,0.04,0.05,0.06,0.07,0.08,0.09,0.1,0.11,0.12,0.13,0.14,0.15,0.16,0.17,0.18,0.19,0.2]
PRRRA=[0,0.01,0.02,0.03,0.04,0.05,0.06,0.07,0.08,0.09,0.1,0.11,0.12,0.13,0.14,0.15,0.16,0.17,0.18,0.19,0.2]
PRAAA=[0,0.01,0.02,0.03,0.04,0.05,0.06,0.07,0.08,0.09,0.1,0.11,0.12,0.13,0.14,0.15,0.16,0.17,0.18,0.19,0.2]
PAARR=[0,0.01,0.02,0.03,0.04,0.05,0.06,0.07,0.08,0.09,0.1,0.11,0.12,0.13,0.14,0.15,0.16,0.17,0.18,0.19,0.2]
PRRAA=[0,0.01,0.02,0.03,0.04,0.05,0.06,0.07,0.08,0.09,0.1,0.11,0.12,0.13,0.14,0.15,0.16,0.17,0.18,0.19,0.2]
id="halfSiblings1_g3-b4-i1"

rule all:
    input:
        expand('data/vcferr/{id}.prarr{prarr}_paara{paara}_prrra{prrra}_praaa{praaa}_paarr{paarr}_prraa{prraa}.vcf.gz',prarr=PRARR, paara=0,prrra=0,praaa=0,paarr=0,prraa=0,id=id),
        expand('data/vcferr/{id}.prarr{prarr}_paara{paara}_prrra{prrra}_praaa{praaa}_paarr{paarr}_prraa{prraa}.vcf.gz.tbi',prarr=PRARR,paara=0,prrra=0,praaa=0,paarr=0,prraa=0,id=id),
        expand('data/akt/{id}.prarr{prarr}_paara{paara}_prrra{prrra}_praaa{praaa}_paarr{paarr}_prraa{prraa}.akt',prarr=PRARR,paara=0,prrra=0,praaa=0,paarr=0,prraa=0,id=id),
        expand('data/vcferr/{id}.prarr{prarr}_paara{paara}_prrra{prrra}_praaa{praaa}_paarr{paarr}_prraa{prraa}.vcf.gz',prarr=0,paara=PAARA,prrra=0,praaa=0,paarr=0,prraa=0,id=id),
        expand('data/vcferr/{id}.prarr{prarr}_paara{paara}_prrra{prrra}_praaa{praaa}_paarr{paarr}_prraa{prraa}.vcf.gz.tbi',prarr=0,paara=PAARA,prrra=0,praaa=0,paarr=0,prraa=0,id=id),
        expand('data/akt/{id}.prarr{prarr}_paara{paara}_prrra{prrra}_praaa{praaa}_paarr{paarr}_prraa{prraa}.akt',prarr=0,paara=PAARA,prrra=0,praaa=0,paarr=0,prraa=0,id=id),
        expand('data/vcferr/{id}.prarr{prarr}_paara{paara}_prrra{prrra}_praaa{praaa}_paarr{paarr}_prraa{prraa}.vcf.gz',prarr=0,paara=0,prrra=PRRRA,praaa=0,paarr=0,prraa=0,id=id),
        expand('data/vcferr/{id}.prarr{prarr}_paara{paara}_prrra{prrra}_praaa{praaa}_paarr{paarr}_prraa{prraa}.vcf.gz.tbi',prarr=0,paara=0,prrra=PRRRA,praaa=0,paarr=0,prraa=0,id=id),
        expand('data/akt/{id}.prarr{prarr}_paara{paara}_prrra{prrra}_praaa{praaa}_paarr{paarr}_prraa{prraa}.akt',prarr=0,paara=0,prrra=PRRRA,praaa=0,paarr=0,prraa=0,id=id),
        expand('data/vcferr/{id}.prarr{prarr}_paara{paara}_prrra{prrra}_praaa{praaa}_paarr{paarr}_prraa{prraa}.vcf.gz',prarr=0,paara=0,prrra=0,praaa=PRAAA,paarr=0,prraa=0,id=id),
        expand('data/vcferr/{id}.prarr{prarr}_paara{paara}_prrra{prrra}_praaa{praaa}_paarr{paarr}_prraa{prraa}.vcf.gz.tbi',prarr=0,paara=0,prrra=0,praaa=PRAAA,paarr=0,prraa=0,id=id),
        expand('data/akt/{id}.prarr{prarr}_paara{paara}_prrra{prrra}_praaa{praaa}_paarr{paarr}_prraa{prraa}.akt',prarr=0,paara=0,prrra=0,praaa=PRAAA,paarr=0,prraa=0,id=id),
        expand('data/vcferr/{id}.prarr{prarr}_paara{paara}_prrra{prrra}_praaa{praaa}_paarr{paarr}_prraa{prraa}.vcf.gz',prarr=0,paara=0,prrra=0,praaa=0,paarr=PAARR,prraa=0,id=id),
        expand('data/vcferr/{id}.prarr{prarr}_paara{paara}_prrra{prrra}_praaa{praaa}_paarr{paarr}_prraa{prraa}.vcf.gz.tbi',prarr=0,paara=0,prrra=0,praaa=0,paarr=PAARR,prraa=0,id=id),
        expand('data/akt/{id}.prarr{prarr}_paara{paara}_prrra{prrra}_praaa{praaa}_paarr{paarr}_prraa{prraa}.akt',prarr=0,paara=0,prrra=0,praaa=0,paarr=PAARR,prraa=0,id=id),
        expand('data/vcferr/{id}.prarr{prarr}_paara{paara}_prrra{prrra}_praaa{praaa}_paarr{paarr}_prraa{prraa}.vcf.gz',prarr=0,paara=0,prrra=0,praaa=0,paarr=0,prraa=PRRAA,id=id),
        expand('data/vcferr/{id}.prarr{prarr}_paara{paara}_prrra{prrra}_praaa{praaa}_paarr{paarr}_prraa{prraa}.vcf.gz.tbi',prarr=0,paara=0,prrra=0,praaa=0,paarr=0,prraa=PRRAA,id=id),
        expand('data/akt/{id}.prarr{prarr}_paara{paara}_prrra{prrra}_praaa{praaa}_paarr{paarr}_prraa{prraa}.akt',prarr=0,paara=0,prrra=0,praaa=0,paarr=0,prraa=PRRAA,id=id)

rule vcferr:
    input: 'data/input/simulation.vcf.gz'
    output: 'data/vcferr/{id}.prarr{prarr}_paara{paara}_prrra{prrra}_praaa{praaa}_paarr{paarr}_prraa{prraa}.vcf.gz'
    shell:
        """ vcferr {input} \
            --sample='{wildcards.id}' \
            --p_rarr={wildcards.prarr} \
            --p_aara={wildcards.paara} \
            --p_rrra={wildcards.prrra} \
            --p_raaa={wildcards.praaa} \
            --p_aarr={wildcards.paarr} \
            --p_rraa={wildcards.prraa} \
            --output_vcf={output} """

rule tabix:
    input: 'data/vcferr/{id}.prarr{prarr}_paara{paara}_prrra{prrra}_praaa{praaa}_paarr{paarr}_prraa{prraa}.vcf.gz'
    output: 'data/vcferr/{id}.prarr{prarr}_paara{paara}_prrra{prrra}_praaa{praaa}_paarr{paarr}_prraa{prraa}.vcf.gz.tbi'
    shell:
        """ tabix {input} """

rule akt:
    input: vcf='data/vcferr/{id}.prarr{prarr}_paara{paara}_prrra{prrra}_praaa{praaa}_paarr{paarr}_prraa{prraa}.vcf.gz',
           tbi='data/vcferr/{id}.prarr{prarr}_paara{paara}_prrra{prrra}_praaa{praaa}_paarr{paarr}_prraa{prraa}.vcf.gz.tbi'
    output: 'data/akt/{id}.prarr{prarr}_paara{paara}_prrra{prrra}_praaa{praaa}_paarr{paarr}_prraa{prraa}.akt'
    shell:
        """ akt kin --force -M1 {input.vcf} > {output} """
```

#### 2.6 R code to generate Figure 1 (`figure1.R`)

```r
library(tidyverse)

if (!file.exists(here::here("figure1.preprocessed.rd"))) {

  message("Preprocessing data...")
  library(skater)

  read_akt2 <- function(fp, varnames = c("prarr", "paara", "prrra", "praaa", "paarr", "prraa")) {
    read_akt(fp) %>%
      mutate(file = basename(fp)) %>%
      mutate(params = gsub("halfSiblings1_g3-b4-i1\\.|\\.akt", "", file)) %>%
      separate(params, into = varnames, sep = "_", remove = FALSE) %>%
      mutate_at(.vars = varnames,  .funs = list(~ as.numeric(gsub("[[:alpha:]]", "", .))))
  }

  tmp_fam <- read_fam("data/input/simulation-everyone.fam")
  tmp_ped <- fam2ped(tmp_fam)

  truth_degrees <-
    ped2kinpair(tmp_ped$ped[[1]]) %>%
    filter(id1 == "halfSiblings1_g3-b4-i1" | id2 == "halfSiblings1_g3-b4-i1") %>%
    mutate(degree = kin2degree(k, max_degree = 3)) %>%
    mutate(degree = ifelse(is.na(degree), "Unrelated", degree)) %>%
    rename(truth_degree = degree, truth_k = k) %>%
    arrange_ids(., id1, id2)

  ## read in akt results
  fps <- list.files("data/akt/", full.names = TRUE)

  res <- map_df(fps, read_akt2)

  ## filter akt results to just pairs with individual sim-ed
  ## join to truth degrees
  res <-
    res %>%
    filter(id1 == "halfSiblings1_g3-b4-i1" | id2 == "halfSiblings1_g3-b4-i1") %>%
    mutate(degree = kin2degree(k, max_degree = 3)) %>%
    mutate(degree = ifelse(is.na(degree), "Unrelated", degree)) %>%
    arrange_ids(id1, id2) %>%
    left_join(truth_degrees, by = c("id1","id2"))

  ## group_by first so we can use the group_keys to get and store param names
  res_grouped <-
    res %>%
    group_by(params)

  ## map the confusion_matrix function over list of akt results
  confmatres <-
    res_grouped %>%
    group_split() %>%
    set_names(unlist(group_keys(res_grouped))) %>%
    map(., ~confusion_matrix(prediction = .x$degree, target = .x$truth_degree))

  warnings()

  ## define a helper to map over conusion matrix list and grab the accuracy
  get_acc <- function(x) {
    x %>%
      pluck("Accuracy")
  }

  ## map over confusion matrix list
  ## use names of list to pull out error params used
  ## misc clean up for params to help with plotting later
  accuracy_dat <-
    map_df(confmatres, get_acc) %>%
    mutate(params = names(confmatres)) %>%
    separate(params,
             into = c("prarr", "paara", "prrra", "praaa", "paarr", "prraa"),
             sep = "_",
             remove = FALSE) %>%
    mutate_at(.vars = c("prarr", "paara", "prrra", "praaa", "paarr", "prraa"),
              .funs = list(~ as.numeric(gsub("[[:alpha:]]", "", .)))) %>%
    mutate(total_error = prarr + paara + prrra + praaa + paarr + prraa) %>%
    mutate(params = gsub("_", ";", params)) %>%
    mutate(params = gsub("p", "", params)) %>%
```

```
    mutate(params = factor(params)) %>%
    mutate(params = fct_reorder(params, total_error))

  accuracy_dat_zeros <- accuracy_dat %>%
    select(Accuracy, prarr:prraa) %>%
    filter(prarr==0 & paara==0 & prrra==0 & praaa==0 & paarr==0 & prraa==0) %>%
    gather(errormode, error_rate, -Accuracy)

  save(accuracy_dat, accuracy_dat_zeros, file=here::here("figure1.preprocessed.rd"))
} else {
  message("Loading preprocessed data...")
  load(here::here("figure1.preprocessed.rd"))
}

accuracy_dat %>%
  select(Accuracy, prarr:prraa) %>%
  gather(errormode, error_rate, -Accuracy) %>%
  filter(error_rate>0) %>%
  bind_rows(accuracy_dat_zeros) %>%
  mutate(`Error mode`=chartr("ar", "AR", errormode)) %>%
  mutate(`Error mode`=`Error mode` %>% factor() %>% fct_reorder(-Accuracy, .fun=mean)) %>%
  ## code to extract incoming genoytpe
  # mutate(ingen = str_extract(`Error mode`, "^.{3}")) %>%
  # mutate(ingen = gsub("p","",ingen)) %>%
  ggplot(aes(error_rate, Accuracy)) +
  geom_point(aes(col=`Error mode`, pch = `Error mode`)) +
  geom_hline(yintercept=unique(accuracy_dat$`Accuracy Guessing`), lty=1, lwd=1.25) +
  geom_line(aes(col=`Error mode`)) +
  scale_x_continuous(breaks = seq(0,0.2, by=0.01)) +
  # facet_wrap(~ingen, ncol = 3)
  labs(x="Error rate", y="Classification accuracy",
       title=NULL,
       subtitle=NULL) +
  theme_bw()

ggsave("figure1.png", width=10, height=6, dpi = 350)

library(geomtextpath)
accuracy_dat %>%
  select(Accuracy, prarr:prraa) %>%
  gather(errormode, error_rate, -Accuracy) %>%
  filter(error_rate>0) %>%
  bind_rows(accuracy_dat_zeros) %>%
  mutate(`Error mode`=chartr("ar", "AR", errormode)) %>%
  mutate(`Error mode`=`Error mode` %>% factor() %>% fct_reorder(-Accuracy, .fun=mean)) %>%
  ggplot(aes(error_rate, Accuracy)) +
  geom_hline(yintercept=unique(accuracy_dat$`Accuracy Guessing`), lty=1, lwd=1.25) +
  geom_textline(aes(label=`Error mode`, group=`Error mode`, col=`Error mode`),
                straight=TRUE, hjust=.15, vjust=.4, fontface=2) +
  scale_x_continuous(breaks = seq(0,0.2, by=0.01)) +
  labs(x="Error rate", y="Classification accuracy",
       title=NULL,
       subtitle=NULL) +
  theme_bw() +
  theme(legend.position="none")

ggsave("figure1.textline.png", width=10, height=6, dpi = 350)
```
